# Genomic evidence for Z-W differentiation in Pacific saury (*Cololabis saira*) and sex-chromosome diversity in Beloniformes

**DOI:** 10.64898/2026.09.08.749792

**Authors:** Keishi Ujiie, Tsubasa Kato, Eitaro Sawayama, Yoji Yamamoto

## Abstract

Genetic sex-determination mechanisms evolve rapidly in teleost fishes and are diverse within Beloniformes. In the present study, the sex-determination system of Pacific saury (*Cololabis saira*) was confirmed using double-digest, restriction-site-associated DNA sequencing (ddRAD-seq) and pooled whole-genome sequencing (Pool-seq), and a DNA marker was developed for inferring genetic sex. DNA sequence analysis detected numerous markers occurring predominantly or exclusively in phenotypic females suggesting a ZZ/ZW sex-determination system. Mapping of ddRAD-seq loci showed that these markers were concentrated within a 3–19 Mb region of chromosome 17 (Chr17). Pool-seq analyses also revealed elevated *F*_ST_, sex-specific differences in sequencing depth, and nucleotide diversity within this region, supporting a ZZ/ZW sex-determination system involving Chr17. A PCR marker designated female-specific 1 (FS1), developed from a female-associated ddRAD sequence unmapped from the male-derived reference genome, was detected in all 43 phenotypic females and none of the 50 phenotypic males collected in 2025. In an independent sample collected in 2026, FS1 was detected in all 19 phenotypic females and one of the 29 phenotypic males, resulting in 97.9% concordance with phenotypic sex. The FS1-positive male may represent a genotypic-phenotypic sex mismatch or reflect recombination between FS1 and the causal sex-determining region; the linkage of FS1 to the W chromosome remains unconfirmed. Chromosome-scale synteny analysis showed that Pacific saury Chr17 and the sex chromosomes of *Hyporhamphus sajori* and *Oryzias latipes* are nonhomologous to one another. These findings reveal substantial sex-chromosome diversity among beloniform fishes and establish a basis for investigating genetic sex determination and genotypic-phenotypic sex concordance in wild Pacific saury.

## Introduction

Genetic sex-determination mechanisms in teleost fishes are remarkably diverse, encompassing variation in master sex-determining genes, sex chromosomes, and heterogametic systems, including male heterogamety (XX/XY) and female heterogamety (ZZ/ZW) (Nagahama et al. 2021; Kitano et al. 2024; Yamamoto and Luckenbach 2024). Rather than being conserved across teleosts, different genes and chromosomes have repeatedly been recruited for sex determination, such that even closely related species may differ in their master sex-determining genes, heterogametic systems, or sex chromosomes (Kitano et al. 2024; Yamamoto and Luckenbach 2024). Comparative analyses of closely related taxa are therefore particularly informative for understanding the evolutionary turnover and diversification of sex-determination systems.

Considerable diversity in sex-determination systems has been documented in several beloniform lineages, particularly in the genera *Oryzias* and *Hyporhamphus*. In the medaka *Oryzias latipes*, male development is initiated by *dmy*, also known as *dmrt1bY*, a Y-linked gene located on chromosome 1 that originated through duplication of the autosomal gene *dmrt1* (Matsuda et al. 2002; Nanda et al. 2002). Other *Oryzias* species possess different master sex-determining genes, including *gsdfY* in *O. luzonensis* and *sox3Y* in *O. dancena*, whereas *amh* has been proposed as a candidate sex-determining gene in *O. eversi* (Myosho et al. 2012; Takehana et al. 2014; Ansai et al. 2022; Kitano et al. 2024). Although male heterogamety occurs in multiple *Oryzias* species, *O. hubbsi* and *O. javanicus* exhibit female heterogamety and possess nonhomologous sex chromosomes, demonstrating that the same heterogametic system can involve different chromosomes (Takehana et al. 2008; Matsuda and Sakaizumi 2016). Similar variation occurs in halfbeaks of the genus *Hyporhamphus*: *H. sajori* has a ZZ/ZW system with an approximately 26-Mb sex-linked region on chromosome 5, whereas *H. intermedius* has an XX/XY system associated with an approximately 140-kb region on chromosome 14 (Xing et al. 2025). Further diversity has been reported in needlefish *Pseudotylosurus microps*, which possess a multiple sex-chromosome system of the X₁X₁X₂X₂/X₁X₂Y type, likely derived from chromosome fusion (Guimarães et al. 2024). This diversity makes Beloniformes a useful group for investigating the evolutionary turnover of sex-determining systems and sex chromosomes.

Beyond their evolutionary significance, identifying sex chromosomes is also important for evaluating how environmental change may affect wild fish populations. Genetic and environmental influences on sex determination are not necessarily mutually exclusive. Even gonochoristic species with well-defined master sex-determining genes can undergo environmentally induced sex reversal when conditions experienced during early development affect sexual differentiation (Devlin and Nagahama 2002). Water temperature is a major environmental factor influencing sexual differentiation in fishes (Hattori et al. 2020). In many species, exposure to elevated temperatures during early development biases sexual differentiation toward males, often through the masculinization of genetic females, although both the magnitude and direction of the response vary among species (Kitano et al. 2024). For example, high temperature induces the development of sex-reversed XX males in *O. latipes* (Hattori et al. 2007). Climate change–driven increases in water temperature may therefore alter phenotypic sex ratios in species whose sexual differentiation is environmentally sensitive (Strüssmann et al. 2010; Geffroy and Wedekind 2020; Lema et al. 2024). In many fish species, environmentally sex-reversed individuals are viable and reproductively functional (Devlin and Nagahama 2002). Their reproduction can affect not only the phenotypic sex ratio of the exposed generation but also the genotypic and phenotypic sex composition of subsequent generations. These effects may have consequences for reproductive success, population dynamics, and population persistence (Schacht et al. 2022). However, biased phenotypic sex ratios in wild populations can also arise from sex-specific differences in migration, survival, and reproductive ecology. Determining whether, and to what extent, sex reversal contributes to population sex-ratio dynamics thus requires individual-level comparisons between genotypic and phenotypic sex, together with an assessment of environmental conditions experienced during sexual differentiation. Reliable molecular markers that identify genotypic sex independently of phenotypic sex are essential for making these comparisons. Characterizing the genetic sex-determination system and its associated genomic regions therefore provides a basis for developing such markers and assessing environmentally induced sex reversal in wild populations.

The Pacific saury *Cololabis saira* (Beloniformes: Scomberesocidae) is a pelagic fish widely distributed in the North Pacific and an important fisheries resource in Japan and other East Asian countries (Fuji et al. 2019). In recent years, its biomass has declined markedly, accompanied by changes in its distribution and migration routes (Hashimoto et al. 2020; Fuji et al. 2023; Nakamura et al. 2024). Although its ecology, migration, reproduction, and population dynamics have been extensively studied, its genetic sex-determination system remains unknown. Recent genomic studies have generated a draft genome assembly accompanied by tissue-specific transcriptomic resources and a male-derived genome assembly comprising 24 chromosome-scale sequences (Nakamura et al. 2024; Sato et al. 2024). However, its sex chromosomes and the extent of sex-linked genomic differentiation have not been identified, and molecular markers for determining genotypic sex are unavailable. This lack of information limits comparative investigations of sex-chromosome evolution within Beloniformes and precludes individual-level comparisons between genotypic and phenotypic sex in Pacific saury, including assessments of potential sex reversal in wild populations. Understanding its reproductive biology, including its sex-determination system, is therefore important not only for addressing these fundamental questions but also for the assessment and sustainable management of this economically important resource.

Here, we compared phenotypic males and females using double-digest restriction-site-associated DNA sequencing (ddRAD-seq) and pooled whole-genome sequencing (Pool-seq) to identify sex-associated genomic regions, characterize sex-specific genomic differentiation, and develop a molecular marker for inferring genotypic sex in Pacific saury. We also compared chromosome-scale synteny among Pacific saury, *H. sajori*, and *O. latipes* to examine sex-chromosome diversity within Beloniformes. By integrating these approaches, we sought to advance understanding of sex-chromosome evolution within Beloniformes and provide a foundation for future studies of genotypic–phenotypic sex concordance and population sex-ratio dynamics in Pacific saury.

## Materials and methods

### Fish Samples and DNA Extraction

Adult Pacific saury were obtained in two different years in Japan. In September 2025, 93 individuals were obtained from Kamaishi, Iwate Prefecture, comprising 43 phenotypic females and 50 phenotypic males. In August 2026, an additional 48 individuals were obtained from Akkeshi, Hokkaido, comprising 19 phenotypic females and 29 phenotypic males. Phenotypic sex was determined by visual inspection of the gonads after dissection. Caudal fin tissue was collected from each individual and preserved in 99.5% ethanol until DNA extraction. The 2025 Kamaishi sample was used for ddRAD-seq and Pool-seq analyses and for the development and initial evaluation of the FS1 marker (see below), whereas the 2026 Akkeshi sample was used for independent validation of the association between FS1 amplification and phenotypic sex.

Approximately 30 mg of fin tissue was digested overnight at 37 °C in 200 µL of TNES-U buffer (Asahida et al. 1996) containing 50 µg of proteinase K. The resulting lysate was treated with 100 µg of RNase A for 5 min at room temperature, and genomic DNA was subsequently purified using homemade SeraPure magnetic beads following the method of Mayjonade et al. (2016). Purified DNA was dissolved in 10 mM Tris-HCl (pH 8.0). DNA integrity was assessed by electrophoresis on a 1% agarose gel, and DNA concentration was measured using a Qubit fluorometer (Thermo Fisher Scientific, USA). DNA samples were adjusted to 50 ng/µL with 10 mM Tris-HCl (pH 8.0) and stored at −20 °C until further analysis.

### ddRAD-seq Library Preparation and Read Processing

Forty-three phenotypic females and 50 phenotypic males were subjected to double-digest restriction-site-associated DNA sequencing (ddRAD-seq). Libraries were prepared following Salas-Lizana and Oono (2018), with modifications based on Maekawa et al. (2023). For each individual, 300 ng of genomic DNA was digested with *Eco*RI-HF and *Mse*I at 37 °C for 4 h, followed by heat inactivation of the restriction enzymes at 65 °C for 10 min. Adapters were then ligated to the digested DNA at 21 °C for 14 h, and the ligase was heat-inactivated at 65 °C for 10 min.

The adapter-ligated DNA was directly amplified using Nextera-based 8-bp dual-index primers (Maekawa et al. 2023). PCR was performed in a total volume of 15 µL containing 3.0 µL of adapter-ligated DNA, 0.2 µM of each index primer, 0.2 mM of each dNTP, 0.3 µL (0.375 U) of PrimeSTAR GXL DNA Polymerase, and 1× PrimeSTAR GXL Buffer (Takara Bio, Japan). Following initial denaturation at 98 °C for 2 min, amplification was performed for 15 cycles of 98 °C for 10 s, 54 °C for 15 s, and 68 °C for 30 s, followed by a final extension at 68 °C for 2 min. An equal volume of the amplified product from each individual was pooled into a single library. Fragments of approximately 450 bp were isolated by double-sided size selection using SPRIselect reagent (Beckman Coulter, Brea, CA, USA). The size distribution and concentration of the resulting library were assessed using a Qsep1 system (BiOptic Inc., Taiwan). The library was sequenced on a NovaSeq X system (Illumina, USA) using 150-bp paired-end sequencing.

Raw reads were demultiplexed by individual based on their 8-bp dual-index sequences. Adapter trimming and quality filtering were performed using fastp v0.25.0 (Chen et al. 2018). Bases with Phred quality scores ≥14 were considered qualified, and reads containing more than 40% unqualified bases, more than 10 ambiguous bases, or fewer than 20 bp after processing were discarded. The resulting high-quality reads were subsequently used to identify sex-linked markers.

### Reference-free Identification of Sex-associated RAD Markers

Sex-associated RAD markers were identified using RADSex v1.2.0 (Feron et al. 2021). Only R1 reads from the quality-filtered ddRAD-seq dataset were used in this reference-free analysis. A marker-depth table was generated using the process command with a minimum read depth of one. Marker-depth statistics for each individual and the frequency distribution of markers across individuals were examined using the depth and freq commands, respectively.

The distribution of markers between phenotypic females and males was evaluated using the distrib command. A marker was considered present in an individual when supported by ≥5 reads. Markers significantly associated with phenotypic sex were extracted using the signif command with the same minimum-depth threshold. Associations between marker presence and phenotypic sex were evaluated using Pearson’s Chi-squared test of independence with Yates’ correction for continuity. Markers with a Bonferroni-corrected *P* value <0.05 were considered significantly associated with phenotypic sex.

Markers showing a female-specific distribution in the 2025 ddRAD-seq dataset were extracted from the marker-depth table using the subset command. These markers were operationally defined as those present at a depth of ≥5 reads in all 43 phenotypic females but absent from all 50 phenotypic males.

### Reference-based Mapping of Sex-associated RAD Markers

RAD markers were aligned to the Pacific saury reference genome assembly (GCF_033807715.1; Sato et al. 2024) using the map command implemented in RADSex (Feron et al. 2021). Only alignments with a minimum mapping quality of 20 were retained. Markers were required to occur at a minimum frequency of 0.1, and a marker was considered present in an individual when supported by ≥5 reads. Marker distributions were compared between phenotypic males and females, specified as groups M and F, respectively.

Associations between marker presence and phenotypic sex were evaluated using Pearson’s Chi-squared test of independence with Yates’ correction for continuity, followed by Bonferroni correction for multiple testing, as implemented in RADSex. The genomic distribution of the resulting sex-association signals was visualized as a Manhattan plot using the ggplot2 package (Wickham 2016) in R v4.5.3 (R Core Team 2026). Bonferroni-corrected *P* values were transformed as −log_10_(*P*), and markers with a corrected *P* value <0.05 were considered significantly associated with phenotypic sex.

### Pool-seq Analysis of Sex-associated Genomic Differentiation

To complement the ddRAD-seq analysis and characterize sex-associated differences in allele frequency, nucleotide diversity, and sequencing depth at higher genomic resolution, pooled whole-genome sequencing (Pool-seq) was performed. Genomic DNA from 30 phenotypic females and 30 phenotypic males randomly selected from the individuals included in the ddRAD-seq analysis was adjusted to 30 ng/µL and pooled separately in equal amounts to generate one female pool and one male pool. Sequencing libraries were constructed separately for the two pools and sequenced on a DNBSEQ platform (MGI Tech, China).

Raw reads were adapter-trimmed and quality-filtered using fastp v0.25.0 (Chen et al. 2018). The filtered paired-end reads were aligned to the Pacific saury reference genome assembly using BWA-MEM v0.7.17-r1188 (Li 2013). The resulting alignments were sorted and indexed using SAMtools v1.21 (Li et al. 2009). Only reads with mapping quality scores ≥20 were retained, and unmapped reads and secondary and supplementary alignments were excluded.

Allele counts were extracted from the filtered BAM files and converted to sync format using grenedalf v0.6.3 (Czech et al. 2024), with a minimum base quality score of 20. Sites were filtered separately for each pool by retaining those with a read depth between 10 and 90 and allele counts ≥2. Only biallelic SNPs with a total minor allele count ≥2 across the two pools were retained for the *F*_ST_ analysis.

Genetic differentiation between the female and male pools was quantified using the unbiased Hudson estimator of *F*_ST_ (Hudson et al. 1992) implemented in grenedalf (Czech et al. 2024). *F*_ST_ was calculated at individual loci and in overlapping 10-kb windows with a step size of 5 kb. Window estimates were averaged across loci that passed all filtering criteria. Genome-wide *F*_ST_ and its 95% confidence interval were additionally estimated using the ANOVA method implemented in the R package poolfstat v3.1.0 (Hivert et al. 2018), with a block-jackknife procedure using blocks of 1,000 SNPs.

Nucleotide diversity (π) was estimated separately for the female and male pools using the diversity command implemented in grenedalf. Estimates were calculated in non-overlapping 10-kb windows using the same pool sizes, mapping- and base-quality thresholds, read-depth thresholds, and minimum within-pool allele count described above. Differences in nucleotide diversity (Δπ) were calculated for each window as

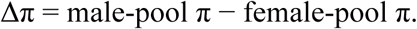

Mean sequencing depth was calculated separately for the female and male pools in non-overlapping 10-kb windows using mosdepth (Pedersen and Quinlan 2018). To account for differences in overall sequencing depth between the two pools, the depth in each window was normalized by the median depth across autosomal windows in the corresponding pool. The candidate sex-associated chromosome and unplaced scaffolds were excluded when calculating autosomal median depth. For pool s and window w, normalized depth was calculated as

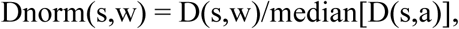

where D(s,w) is the mean sequencing depth in window w and a represents the autosomal windows included in the normalization. The sex-specific depth ( Δ depth) difference for each window was then calculated as

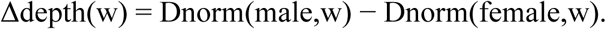

Positive Δdepth values therefore indicate higher normalized sequencing depth in the male pool than in the female pool. Normalized depth profiles and depth differences were visualized along the candidate sex-associated chromosome.

Genome-wide patterns of *F*_ST_ and nucleotide diversity, together with normalized sequencing-depth profiles along the candidate sex-associated chromosome, were visualized in R v4.5.3 using ggplot2 v4.0.3.

### Exploratory Analysis of Sex-associated Structural Variants

Structural variants were identified separately in the female and male Pool-seq datasets using the short-read workflow implemented in DELLY v1.2.6 (Rausch et al. 2012). Deletions, duplications, and inversions were initially called from the filtered BAM file of each pool using the Pacific saury reference genome assembly. Variant sites detected in either pool were merged to generate a common set of candidate structural variants, which was subsequently genotyped separately in the female and male pools using DELLY. The resulting BCF files were merged using BCFtools v1.21 (Danecek et al. 2021).

For each structural variant, reference-supporting read counts were calculated as the sum of paired-end and split-read reference counts, whereas variant-supporting read counts were calculated as the sum of paired-end and split-read variant counts. Variants were retained when the total number of supporting reads was between 10 and 200 in each pool. Differences in the proportions of reference- and variant-supporting reads between the female and male pools were evaluated using Fisher’s exact test. The resulting *P* values were corrected for multiple testing using the Benjamini–Hochberg procedure.

Structural variants were assigned to non-overlapping 10-kb windows based on their genomic start positions. For each window, the maximum −log_10_(false-discovery-rate-adjusted *P* value) among the variants within that window was used for visualization. Windows containing structural variants with an adjusted *P* value <0.001 were considered candidates showing sex-biased read support. Because only one pool was analyzed for each phenotypic sex, this analysis was considered exploratory and did not estimate biological variation among replicate pools.

### Development and Evaluation of the Female-associated Molecular Marker FS1

Four female-associated ddRAD sequences meeting the marker-selection criteria described above were selected for PCR marker development. Primer pairs were designed and assessed for sequence specificity using NCBI Primer-BLAST (Ye et al. 2012). The four candidate female-specific (FS) primer pairs were initially screened using genomic DNA from 12 phenotypic females and 12 phenotypic males from the sample collected in 2025. Three of the four candidate primer pairs produced amplification products in both sexes, whereas the primer pair designated “FS1” produced an amplification product only in phenotypic females and was therefore selected for further evaluation. The primer sequences and PCR product sizes for the three candidate markers that amplified in both sexes are provided in Supplementary Table S1.

For molecular sex identification, a multiplex PCR assay was developed by combining the FS1 primer pair with a primer pair targeting β-actin as an internal positive control. The β-actin primers were designed based on the predicted Pacific saury β-actin transcript available in the NCBI database (accession no. XM_061719902). The primer sequences were as follows: FS1 forward, 5′-CCCAACATACTCCAGGTGCT-3′; FS1 reverse, 5′-CCAAGACACTAATGGCGCTT-3′; β-actin forward, 5′-CCCAGGCATCAGGGTGTAAT-3′; and β-actin reverse, 5′-AGAGGCAGCAGTTCCCATTT-3′. The expected product sizes were 120 bp for FS1 and 585 bp for β-actin.

Multiplex PCR was performed in a total volume of 10 µL containing 6.35 µL of distilled water, 1.0 µL of 10× Ex Taq Buffer, 0.8 µL of dNTP mixture, 0.1 µL each of the forward and reverse β-actin primers, 0.3 µL each of the forward and reverse FS1 primers, 0.05 µL (0.25 U) of Ex Taq DNA Polymerase (5 U/µL; Takara Bio), and 1.0 µL of template DNA. All primer stock solutions were 10 µM, resulting in final concentrations of 0.1 µM for each β-actin primer and 0.3 µM for each FS1 primer. Following initial denaturation at 98 °C for 2 min, amplification was performed for 28 cycles of 98 °C for 10 s, 66 °C for 30 s, and 72 °C for 30 s, followed by a final extension at 72 °C for 2 min. Amplification products were separated by electrophoresis on a 1.0% agarose gel and visualized alongside a 100-bp DNA ladder.

The association between FS1 amplification and phenotypic sex was initially evaluated in all 93 individuals from the sample used for ddRAD-seq, comprising 43 phenotypic females and 50 phenotypic males. Representative amplification patterns from 12 phenotypic females and 12 phenotypic males were visualized by agarose-gel electrophoresis. This association was independently evaluated in an additional 48 individuals from the sample collected in 2026, comprising 19 phenotypic females and 29 phenotypic males. An individual was scored as FS1-positive when both the 120-bp FS1 product and the 585-bp β-actin internal-control product were detected and as FS1-negative when only the β-actin product was detected.

### Comparative Chromosome-scale Synteny Analysis

Chromosome-scale genome assemblies of Pacific saury, *H. sajori*, and *O. latipes* were analyzed to compare syntenic relationships among the three species. Genome completeness and conserved ortholog content were assessed using BUSCO v5.5.0 in genome mode with the actinopterygii_odb10 (Manni et al. 2021). The genome assemblies used were Pacific saury (GCF_033807715.1), *H. sajori* (GCA_050306495.1), and *O. latipes* (GCF_053564925.1).

Complete single-copy BUSCO genes shared among the three species were identified based on their BUSCO identifiers. For each shared ortholog, its chromosome assignment and genomic start and end positions were extracted from the BUSCO full tables. BUSCO genes that were duplicated, fragmented, missing, located on unplaced scaffolds, or not represented exactly once in each species were excluded. Chromosome lengths were obtained from the FASTA index files generated using SAMtools.

Corresponding BUSCO genes were linked between *O. latipes* and Pacific saury and between Pacific saury and *H. sajori*. Syntenic relationships were visualized using NGenomeSyn v1.43 (He et al. 2023), with Pacific saury positioned as the central reference species. Chromosome order in the upper and lower rows was optimized to reduce link crossings, while the original chromosome identifiers and numbering of each genome assembly were retained.

In addition, protein sequences of *amh*, *amhr2*, and *dmrt1* were used as queries in TBLASTN searches against the male-derived Pacific saury genome assembly (GCF_033807715.1). Query species, protein accession numbers, and genomic coordinates of the identified homologs are provided in Supplementary Table S2.

## Results

### Reference-free Identification of Sex-associated RAD Markers

RADSex analysis revealed a pronounced excess of markers occurring predominantly or exclusively in phenotypic females (Fig. 1). Numerous marker distributions showing strong female bias were significantly associated with phenotypic sex after Bonferroni correction. In particular, four markers were detected at a depth of ≥5 reads in all 43 phenotypic females but were completely absent from all 50 phenotypic males. In contrast, no comparable set of markers showing complete male-specific occurrence was detected. This asymmetric distribution of sex-associated RAD markers was consistent with female heterogamety (ZZ/ZW).

**Fig. 1.**
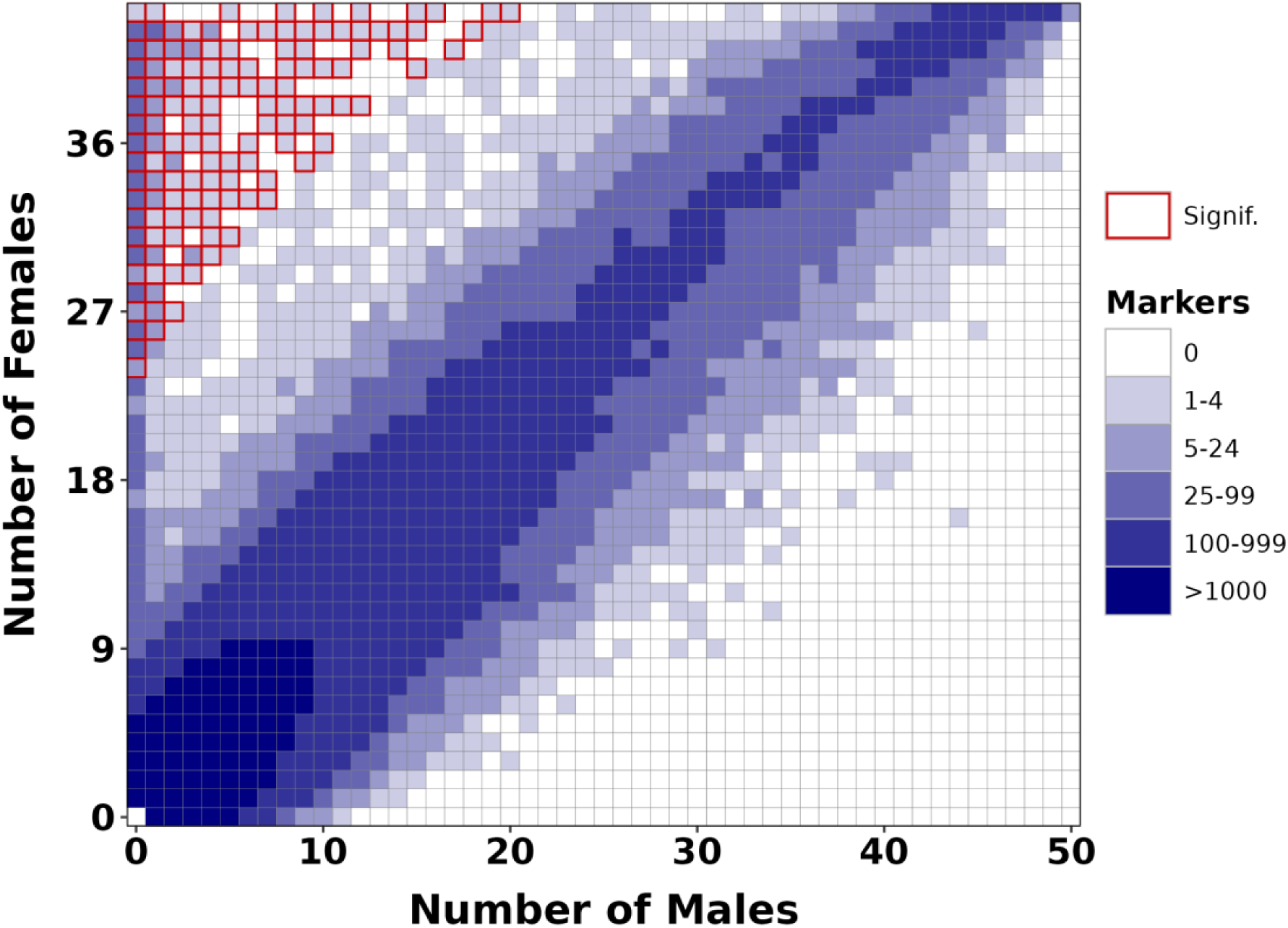
Distribution of ddRAD markers between phenotypic males and females of Pacific saury (*Cololabis saira*). Distribution of RAD markers according to the numbers of phenotypic males and females in which each marker was detected. The color scale indicates the number of markers showing each distribution pattern. Red-outlined cells indicate distributions significantly associated with phenotypic sex based on Chi-squared tests with Bonferroni correction. A marker was considered present in an individual at a sequencing depth of ≥5 reads.

### Genomic Localization of Sex-associated RAD Markers

Reference-based mapping showed that sex-associated RAD markers were strongly concentrated on Chr17 (CM066558.1), whereas relatively few significant markers were distributed across the other chromosomes (Fig. 2a). Within Chr17, significant markers were broadly distributed between approximately 3 and 19 Mb, with many markers showing highly significant associations with phenotypic sex throughout this interval (Fig. 2b). Thus, the reference-free and reference-based RADSex analyses consistently identified Chr17 as the principal chromosome associated with sex.

**Fig. 2.**
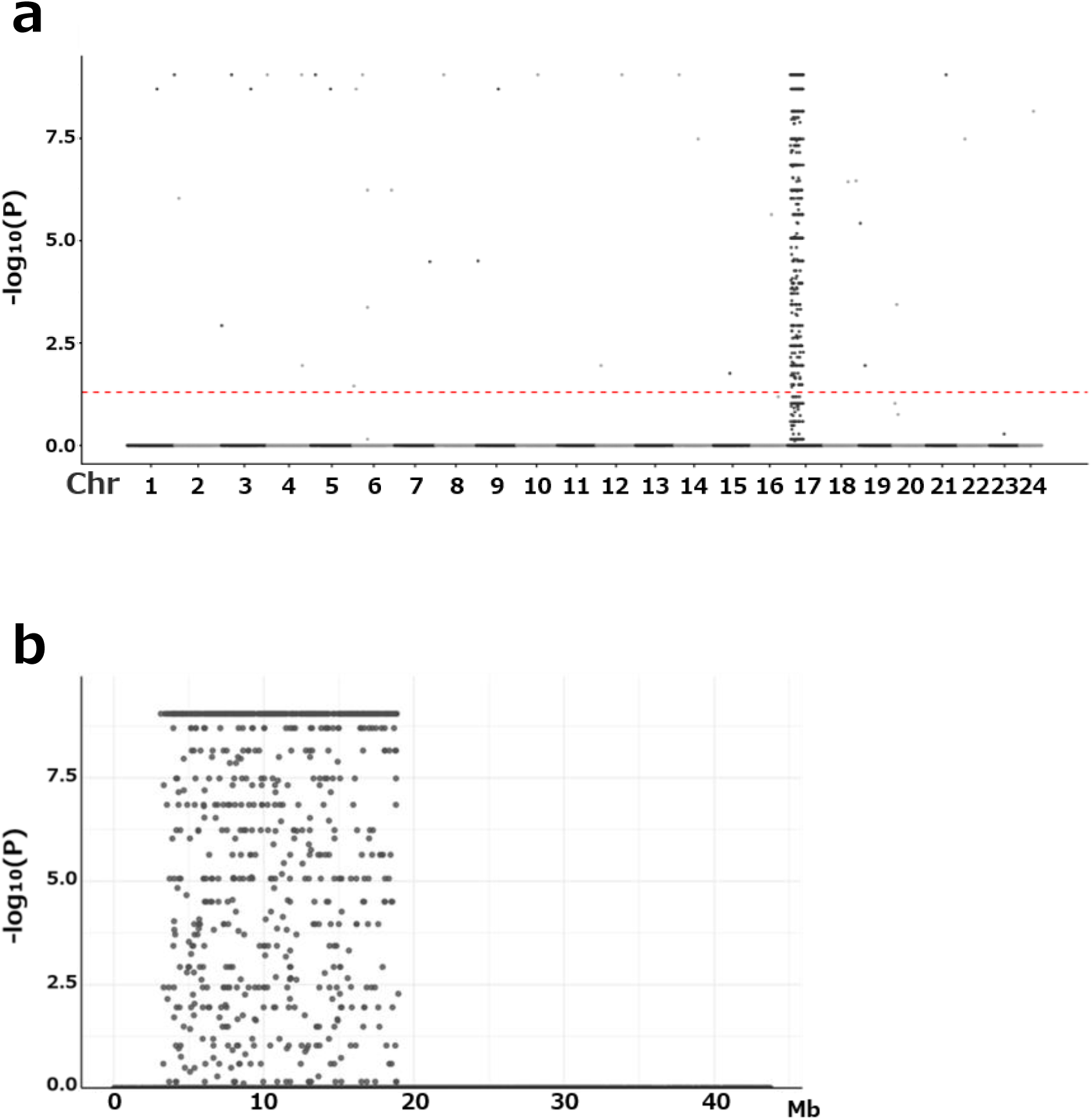
Genomic distribution of sex-associated ddRAD markers in Pacific saury (*Cololabis saira*). **a** Genome-wide distribution of sex-association significance for RAD markers mapped to the male-derived Pacific saury reference genome. Chromosomes are shown in numerical order along the x-axis, and the y-axis represents − log_10_(Bonferroni-corrected *P* value). The red dashed line indicates the significance threshold corresponding to a Bonferroni-corrected *P* value of 0.05. **b** Enlarged view of sex-associated RAD markers mapped to chromosome 17 (GenBank accession CM066558.1). Marker positions are shown along the x-axis, and the y-axis represents −log_10_(Bonferroni-corrected *P* value).

### Pool-seq Analysis of Sex-associated Genomic Differentiation

Pool-seq analysis identified pronounced genomic differentiation between the female and male pools on Chr17. Windowed *F*_ST_ values were elevated across an approximately 16-Mb interval extending from around 3 to 19 Mb, whereas values outside this region were generally close to zero (Fig. 3a). Within the differentiated region, most 10-kb windows showed *F*_ST_ values of approximately 0.15–0.45, with a maximum value exceeding 0.6. The genome-wide *F*_ST_ estimated using poolfstat was 0.0061 (95% CI: −0.0034 to 0.0206).

**Fig. 3.**
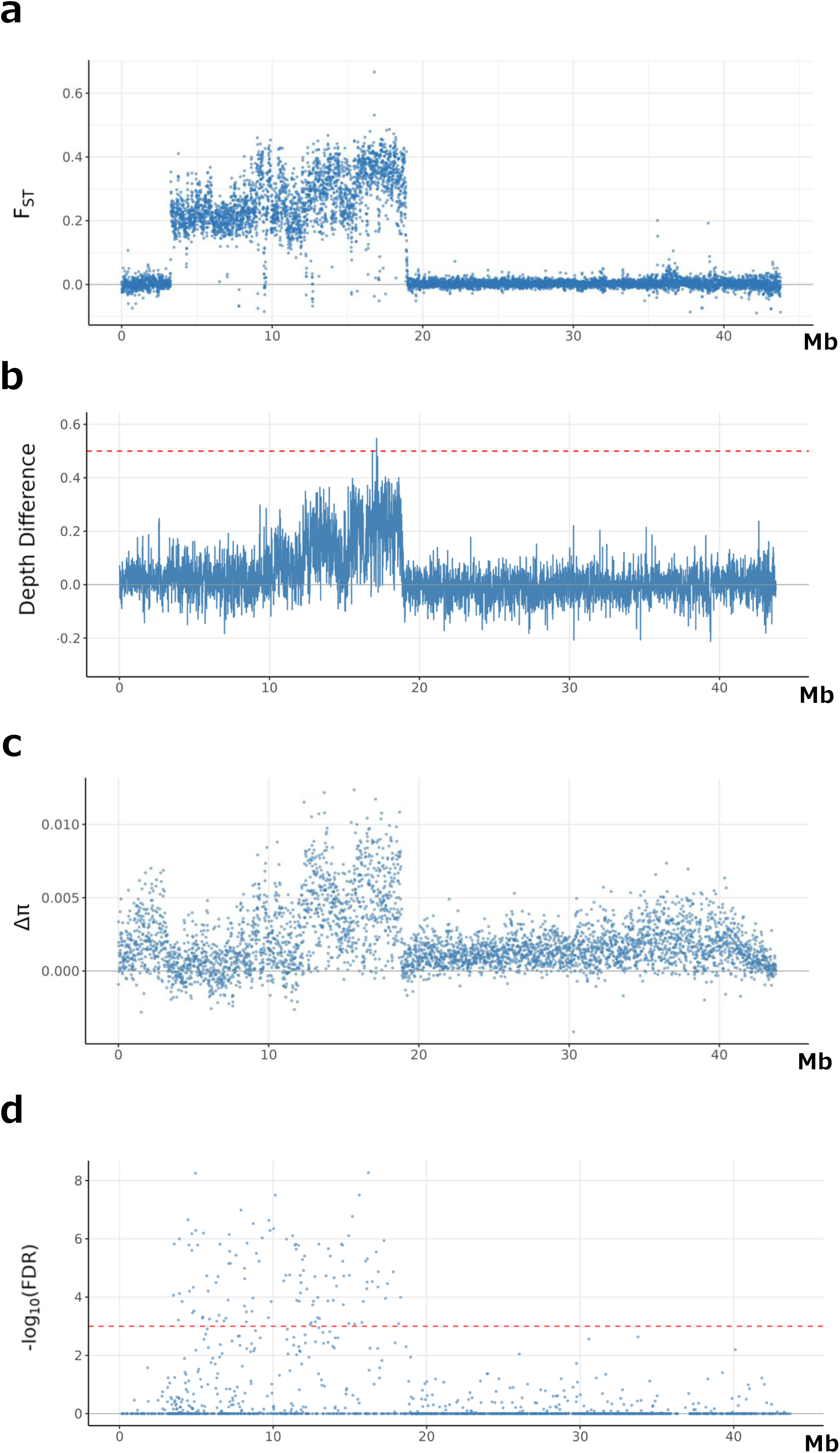
Sex-associated patterns of genomic differentiation, sequencing depth, nucleotide diversity, and deletion-like signals along chromosome 17 of Pacific saury (*Cololabis saira*). **a** *F*_ST_ between the phenotypic female and male pools, calculated using the unbiased Hudson estimator in overlapping 10-kb windows with a 5-kb step size. **b** Difference in normalized sequencing depth between the male and female pools, calculated as normalized male depth minus normalized female depth in non-overlapping 10-kb windows. Sequencing depth in each pool was normalized by the median depth across autosomal windows in that pool. Positive values indicate higher normalized depth in males than in females. The solid horizontal line at zero indicates no difference between the pools. The red dashed line at 0.5 indicates the expected difference when female sequencing depth is half that of males, as predicted for a Z-linked region lacking corresponding mappable W-derived sequences. **c** Difference in nucleotide diversity (π) between the male and female pools, calculated as male-pool π minus female-pool π in non-overlapping 10-kb windows. Positive values indicate higher nucleotide diversity in males than in females. **d** Exploratory distribution of deletion-like signals identified using DELLY. Each point represents a 10-kb window and is plotted according to the maximum −log_10_(Benjamini– Hochberg-adjusted *P* value) among candidate deletions within that window. The red dashed line indicates a Benjamini–Hochberg-adjusted *P* value of 0.001, with points above the line representing adjusted *P* values < 0.001. Chromosome coordinates refer to CM066558.1.

Normalized sequencing depth also differed between the sexes within Chr17. The normalized depth of the male pool was consistently higher than that of the female pool across much of the sex-associated interval, particularly between approximately 10 and 19 Mb (Fig. 3b). The Δdepth reached approximately 0.5 in some windows, whereas differences were generally centered near zero across the remainder of the chromosome. Because the reference genome was derived from a male, the reduced female-pool depth in this region was consistent with substantial sequence differentiation between the Z- and W-linked regions.

Nucleotide diversity differed similarly between the two pools. The Δπ was predominantly positive within the sex-associated region and was especially pronounced between approximately 12 and 19 Mb (Fig. 3c). This pattern indicated lower nucleotide diversity in the female pool than in the male pool across much of the differentiated interval.

Exploratory structural-variant analysis further detected an accumulation of deletions showing sex-biased read support within the same region of Chr17 (Fig. 3d). Most 10-kb windows exceeding the FDR threshold were located between approximately 3 and 18 Mb, whereas few such signals were detected across the remainder of the chromosome. Thus, *F*_ST_, normalized sequencing depth, nucleotide diversity, and deletion-associated read support all identified an overlapping region of pronounced differentiation between the female and male pools.

### Development and Validation of the Female-associated Marker FS1

Four female-associated ddRAD sequences were initially selected for PCR marker development. Three candidate primer pairs produced amplification products in both phenotypic females and males and were therefore excluded from further analysis (Supplementary Table S1). In contrast, during the initial screening, the FS1 primer pair produced the expected 120-bp amplification product in phenotypic females but not in phenotypic males, whereas the 585-bp β-actin internal-control product was successfully amplified in both sexes.

FS1 amplification was examined in all 93 individuals from the sample collected in 2025 (Table 1). The FS1 product was detected in all 43 phenotypic females and was absent from all 50 phenotypic males. These results confirmed that the female-associated ddRAD-seq pattern was reproducible using the FS1 PCR assay. Representative PCR patterns from 12 phenotypic females and 12 phenotypic males are shown in Fig. 4.

**Fig. 4.**
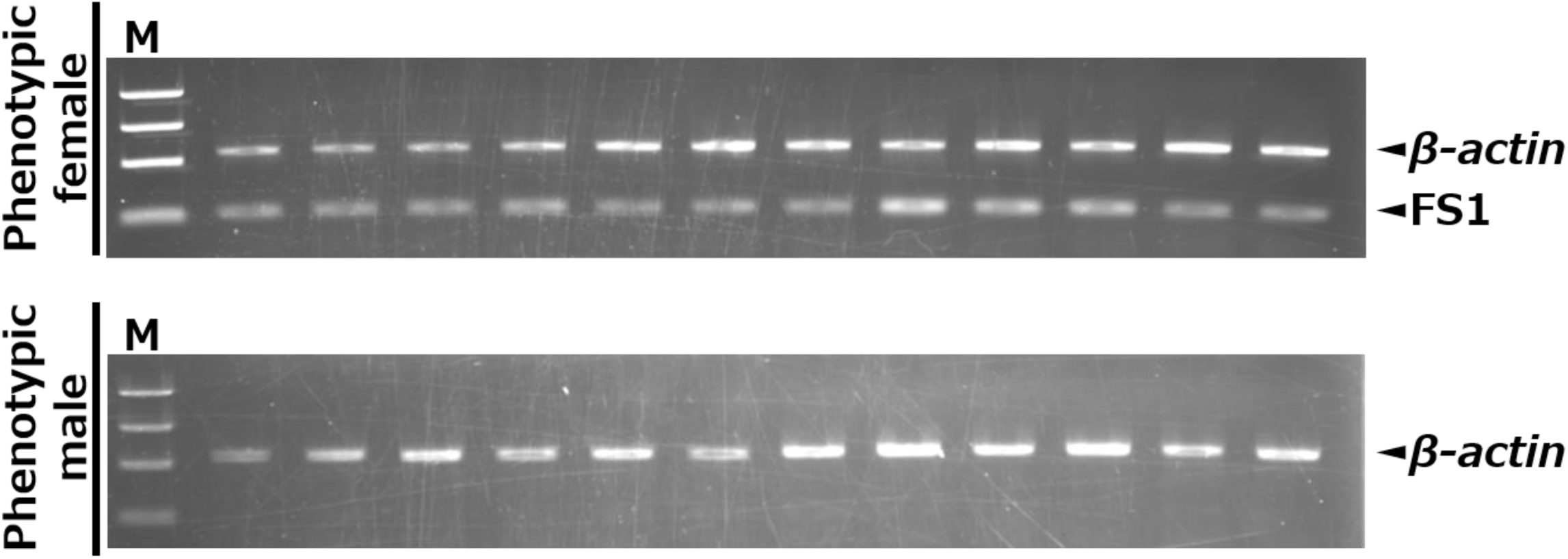
Multiplex PCR amplification of the female-associated marker FS1 in Pacific saury (*Cololabis saira*). Representative amplification patterns are shown for 12 phenotypic females and 12 phenotypic males from the 2025 Kamaishi sample. FS1 produced the expected 120-bp product in phenotypic females but not in phenotypic males, whereas the 585-bp β-actin internal-control product was amplified in all individuals. M, 100-bp DNA ladder.

**Table 1.** Association between phenotypic sex and FS1 amplification in Pacific saury sampled in 2025 and 2026.

| Sampling year<br>(landing location) | Phenotypic sex | FS1-negative<br>(putative ZZ)<br>n (%) | FS1-positive<br>(putative ZW)<br>n (%) | Total<br>n (%) |
| --- | --- | --- | --- | --- |
| 2025 (Kamaishi) | Male | 50 (100) | 0 (0) | 50 (53.8) |
|  | Female | 0 (0) | 43 (100) | 43 (46.2) |
|  | Total | 50 (53.8) | 43 (46.2) | 93 (100) |
| 2026 (Akkeshi) | Male | 28 (100) | 1 (5.0) | 29 (60.4) |
|  | Female | 0 (0) | 19 (95.0) | 19 (39.6) |
|  | Total | 28 (58.3) | 20 (41.7) | 48 (100) |

Phenotypic sex was determined by visual inspection of the gonads after dissection (testis = male; ovary = female). Putative genotypic sex was inferred from FS1 amplification: FS1-negative individuals were classified as putative ZZ, whereas FS1-positive individuals were classified as putative ZW. Percentages in the FS1-negative and FS1-positive columns were calculated relative to the total number of individuals in the corresponding putative genotypic-sex class. Percentages in the total rows and total column were calculated relative to the total number of individuals examined in each annual sample.

The association between FS1 amplification and phenotypic sex was independently evaluated in 48 individuals from the sample in 2026 (Table 1). FS1 was detected in all 19 phenotypic females and in one of the 29 phenotypic males, whereas the remaining 28 phenotypic males were FS1-negative. Thus, the FS1 amplification pattern was concordant with phenotypic sex in 47 of the 48 individuals (97.9%). Accordingly, one of the 20 putative ZW individuals (5.0%) in the 2026 sample exhibited a male gonadal phenotype.

### Comparative Chromosome-scale Synteny among Beloniform Fishes

A total of 3,434 complete single-copy BUSCO orthologs shared among Pacific saury, *H. sajori*, and *O. latipes* were used to compare chromosome-scale synteny (Fig. 5). Most orthologous links connected each Pacific saury chromosome predominantly to a single chromosome in each of the other two species, indicating broad conservation of chromosome-scale macrosynteny, although intrachromosomal gene order varied among species.

**Fig. 5.**
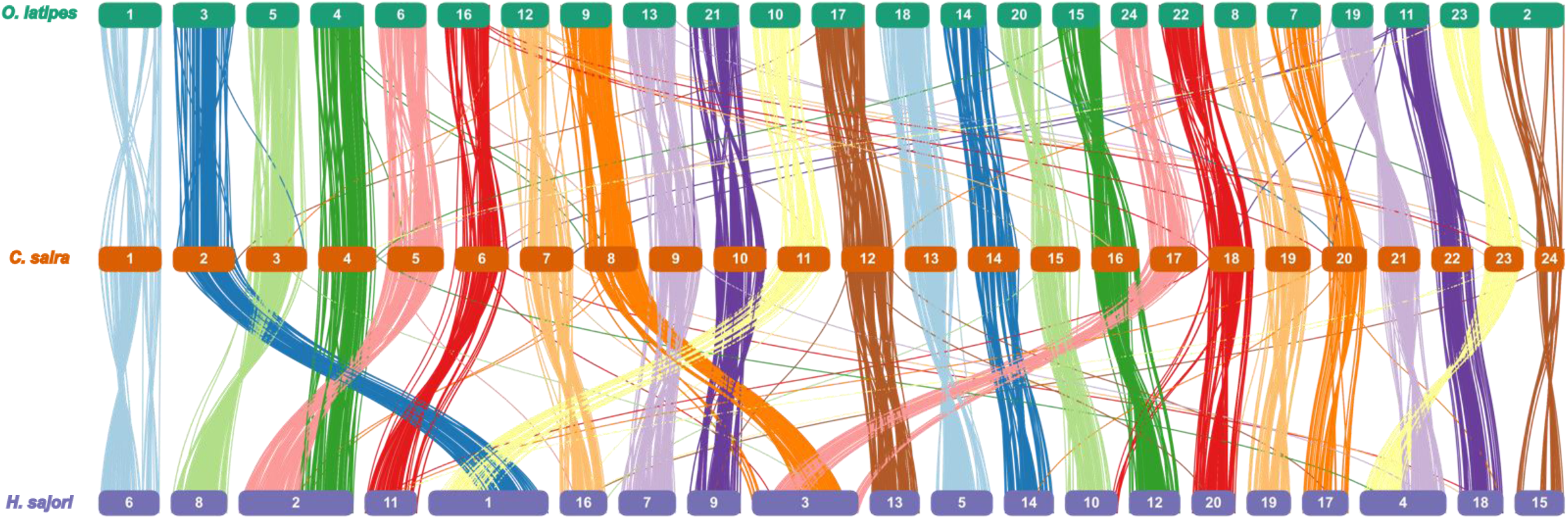
Comparative chromosome-scale synteny among Pacific saury (*Cololabis saira*), halfbeak (*Hyporhamphus sajori*), and medaka (*Oryzias latipes*). Chromosome-scale synteny was assessed using 3,434 complete single-copy BUSCO orthologs shared among the three species. Each line connects the chromosomal positions of the same ortholog in two species. Chromosomes of *O. latipes*, Pacific saury (*C. saira*), and *H. sajori* are shown in the upper, middle, and lower rows, respectively. The sex-associated chromosome of Pacific saury (Chr17) and the sex chromosomes of *H. sajori* (Chr5) and *O. latipes* (Chr1) are nonhomologous to one another.

The sex-associated chromosomes of the three species are nonhomologous to one another. Specifically, Pacific saury Chr17 was nonhomologous to the reported sex chromosomes of both *H. sajori* and *O. latipes*, which were also nonhomologous to each other.

TBLASTN searches identified *amh*, *amhr2*, and *dmrt1* homologs on Chr13, Chr12, and Chr9, respectively; none was located within the Chr17 sex-associated region (Supplementary Table S2).

## Discussion

In this study, genome-wide comparisons between females and males using ddRAD-seq and Pool-seq identified an extensive sex-associated region spanning approximately 3–19 Mb on Chr17 of Pacific saury. Female-biased markers detected by ddRAD-seq were concentrated within this region, where Pool-seq analyses also revealed sex-specific differences in *F*_ST_, sequencing depth, and nucleotide diversity. Furthermore, FS1, a PCR marker developed from a female-associated ddRAD sequence, reproduced this pattern in the sample collected in 2025 and showed 97.9% concordance with phenotypic sex in the independently examined 2026 sample. Taken together, these findings provide genome-wide evidence supporting a ZZ/ZW sex-determination system involving Chr17 in Pacific saury. Chromosome-scale synteny analysis further showed that the sex-associated chromosomes of Pacific saury, *H. sajori*, and *O. latipes* are nonhomologous to one another.

The distribution of sex-associated signals across an approximately 16-Mb region of Chr17 in Pacific saury is consistent with extensive differentiation between the Z and W chromosomes and raises the possibility that recombination is suppressed across a substantial portion of this region. In *H. sajori*, another beloniform fish with a ZZ/ZW system, an approximately 26-Mb region on Chr5 exhibits high sex-specific *F*_ST_, numerous heterozygous SNPs in females, and extensive linkage disequilibrium, supporting its identification as a fully sex-linked region with suppressed recombination (Xing et al. 2025). In Pacific saury, however, recombination rates and linkage disequilibrium were not directly evaluated; therefore, the presence and extent of recombination suppression remain to be determined. *F*_ST_ decreased sharply at approximately 19 Mb, but whether this position represents the boundary of the sex-associated region requires further examination using change-point or linkage analyses.

The Pacific saury chromosome assembly was derived from a male individual (Sato et al. 2024) and, under a ZZ/ZW system, is expected to primarily represent the Z chromosome and lack W-specific sequences. When female reads are mapped to a male-derived reference genome, female coverage is expected to approach half that of males where the corresponding W-linked sequences are absent or substantially diverged (Xing et al. 2025). Consistent with this expectation, female sequencing depth was lower within the sex-associated region of Chr17 and approached half the male depth in some intervals. An exploratory structural-variant analysis using DELLY also revealed a concentration of deletion-like signals with differential read support between the female and male pools. The magnitude of Z–W differentiation was not uniform across the approximately 3–19 Mb interval. Elevated *F*_ST_ extended broadly across this region, whereas the reduction in female sequencing depth was particularly pronounced between approximately 10 and 19 Mb and female nucleotide diversity was most strongly reduced between approximately 12 and 19 Mb. These patterns suggest a mosaic of regions with retained Z–W homology and regions with substantial sequence or structural differentiation, with stronger differentiation-related signals toward the distal portion of the sex-associated region. The reduction in female nucleotide diversity likely reflects the lower representation of Z-derived sequences and reduced mapping of divergent W-derived reads rather than necessarily indicating degeneration of the W chromosome itself. Together, the concordant changes in *F*_ST_, sequencing depth, nucleotide diversity, and structural-variant signals support extensive but spatially heterogeneous Z–W differentiation across the approximately 3–19 Mb region of Chr17. Whether the depth reductions reflect W-linked deletions or extensive sequence divergence cannot be resolved using the male-derived reference genome alone. Likewise, identification of the causal sex-determining gene will require characterization of W-specific sequences and functional analyses and remains an important subject for future study.

The differences among the sex-associated chromosomes of Pacific saury, *H. sajori*, and *O. latipes* may extend to the genes associated with sex determination. In *O. latipes*, the Y-linked *dmy* gene on Chr1, which originated through duplication of *dmrt1*, functions as the master sex-determining gene (Matsuda et al. 2002; Nanda et al. 2002). In *H. sajori*, the Chr5 sex-linked region contains Z- and W-linked copies of *amhr2* that differ at five predicted amino acid residues, making *amhr2* a candidate sex-determining gene (Xing et al. 2025). In *H. intermedius*, *amh* is located within an approximately 140-kb Y-linked region on Chr14, with two strongly sex-associated variants occurring upstream of the gene (Xing et al. 2025). Thus, different members of the TGF-β signaling pathway have been proposed as candidate sex-determining genes in these two *Hyporhamphus* species. In Pacific saury, however, the annotated *amh*, *amhr2*, and *dmrt1* homologs occur on Chr13, Chr12, and Chr9, respectively, rather than within the Chr17 sex-associated region. These annotated copies are therefore unlikely to represent the Chr17-linked master sex-determining factor. Nevertheless, the possibility remains that a W-specific duplicated gene or highly diverged homolog is absent from the male-derived reference genome. Identifying the master sex-determining factor will therefore require a female-derived genome assembly, together with expression and functional analyses of candidate genes during early sexual differentiation.

The occurrence of ZZ/ZW systems involving nonhomologous chromosomes in Pacific saury and *H. sajori* indicates that female heterogamety is associated with different chromosomes in the two species. Pacific saury and *H. sajori* occupy distinct phylogenetic branches within Beloniformes (Setiamarga et al. 2008). Phylogenetic and divergence analyses further indicate that the Chr5-linked ZZ/ZW system of *H. sajori* arose after its divergence from *H. intermedius*, which possesses a Chr14-linked XX/XY system (Xing et al. 2025). Similarly, *Oryzias hubbsi* and *O. javanicus* both possess ZZ/ZW systems, but their sex chromosomes correspond to different linkage groups (Takehana et al. 2008; Matsuda and Sakaizumi 2016). Sex-chromosome turnover may result from the acquisition of a novel sex-determining factor, translocation of an existing sex-determining factor, or genomic rearrangement involving a sex-determining region (Herpin et al. 2021; Kitano et al. 2024; Xing et al. 2025). However, because the ancestral sex-determination system and sex chromosomes of Pacific saury and *H. sajori* remain unknown, it is not possible to determine whether female heterogamety was retained while the sex chromosome changed or evolved independently in the two lineages. Nevertheless, the involvement of different chromosomes in sex determination among Pacific saury, *H. sajori*, and other beloniform fishes demonstrates the substantial evolutionary diversity of sex chromosomes within this lineage.

The FS1 marker was developed from a female-associated ddRAD sequence that did not map to the male-derived reference genome, consistent with its possible linkage to the W chromosome. In the same 93 individuals used for ddRAD-seq, FS1 was detected in all 43 phenotypic females and was absent from all 50 phenotypic males, confirming that the female-associated ddRAD pattern was reproducible using the PCR assay. In the independently examined 2026 sample, FS1 was detected in all 19 phenotypic females and was absent from 28 of the 29 phenotypic males, resulting in 97.9% concordance with phenotypic sex. Nevertheless, the chromosomal location and inheritance pattern of FS1 have not been directly determined. Physical mapping to the W chromosome and family-based analyses of its maternal inheritance and cosegregation with sex will therefore be required to establish FS1 as a genetic sex marker. One FS1-positive phenotypic male was detected in the 2026 sample landed at Akkeshi, Hokkaido, whereas no discordant individuals were detected in the 2025 sample landed at Kamaishi, Iwate. If FS1 is linked to the W chromosome, this individual could represent a putative ZW male exhibiting a mismatch between genotypic and phenotypic sex. Alternatively, recombination between the FS1 locus and the causal sex-determining region could have uncoupled the marker from genotypic sex. The present data do not distinguish between these explanations and therefore do not demonstrate sex reversal. Genotyping with additional markers spanning the sex-associated region and family-based segregation analysis will be required to determine the genotypic sex of this individual. A population-genomic analysis of Pacific saury from eight North Pacific locations reported very low genome-wide differentiation among locations (*F*_ST_ < 0.005; Liu et al. 2024). Although this low differentiation suggests that large-scale population structure may not strongly limit the transferability of FS1, its reliability as a genetic sex marker will require independent validation in samples collected across broader geographic areas, years, and cohorts.

Beyond its application to genetic sex identification, FS1 may provide a basis for investigating how environmental variation affects sexual differentiation and population sex ratios in wild Pacific saury. Environmentally influenced sex determination or sex reversal has been documented in *O. latipes* and several atheriniform fishes, although phylogenetic relatedness alone does not demonstrate that Pacific saury shares this environmental sensitivity (Hattori et al. 2007; Yamamoto et al. 2014; Miyoshi et al. 2020; Strüssmann et al. 2021). Once its W linkage and reliability have been established, comparing genotypic sex inferred from FS1 with gonadal phenotype, hatch timing inferred from daily otolith increments, and environmental temperature data could provide a means of testing whether genotypic–phenotypic sex mismatches are associated with thermal conditions experienced during early development (Miyoshi et al. 2020; Fujimoto et al. 2026). Because sex-reversed individuals can be reproductively functional (Devlin and Nagahama 2002), such analyses could ultimately clarify how environmental variation affects population sex ratios across generations.

In conclusion, this study provides genome-wide evidence supporting a ZZ/ZW sex-determination system involving Chr17 in Pacific saury. Chromosome-scale synteny analysis showed that different chromosomes are associated with sex determination among beloniform fishes, highlighting the evolutionary diversity of sex chromosomes within this lineage. FS1 reproduced the female-associated ddRAD pattern and showed 97.9% concordance with phenotypic sex in the independently examined sample collected in 2026, supporting its potential as a genetic sex marker. However, the physical linkage of FS1 to the W chromosome and its inheritance pattern remain to be confirmed through family-based segregation analysis. The genetic sex of the FS1-positive male from the sample collected in 2026 should also be evaluated using additional sex-linked markers. A female-derived genome assembly will be essential for characterizing the W chromosome and identifying the master sex-determining factor. Following confirmation of its W linkage and broader validation across populations, FS1 could enable surveys of genotypic–phenotypic sex concordance in wild populations, providing a basis for evaluating how environmental variation may affect population sex ratios, with implications for the sustainable management of Pacific saury.

### Data availability

The raw ddRAD-seq and Pool-seq reads generated in this study have been deposited in the DDBJ Sequence Read Archive (DRA) and will be made publicly available upon acceptance of the peer-reviewed article. All other data supporting the findings of this study are included in the article and its supplementary information or are available from the corresponding author upon reasonable request.

## Acknowledgments

We thank Hamako Suisan, Kamaishi, Iwate, Japan, for providing the Pacific saury samples, and Associate Professor Kensuke Ichida (Iwate University) for his assistance in shipping them. We also thank Associate Professor Takashi Ichikawa (Tokyo University of Agriculture) and Tomoe Ichikawa for their assistance in obtaining and shipping the samples from Akkeshi, Hokkaido. We are grateful to Dr. Adam Luckenbach of the Northwest Fisheries Science Center, National Oceanic and Atmospheric Administration (NOAA), for his valuable comments on the manuscript and assistance with English-language editing.

## Declarations Conflicts of interest

The authors declare that they have no competing interests.

## Ethics approval

All experimental procedures involving fish were conducted in accordance with the guidelines for animal experimentation of Tokyo University of Marine Science and Technology and with relevant domestic and international guidelines for the ethical treatment of animals.

## Contributions

KU and TK contributed to the methodology, performed the experiments, curated and analyzed the data, prepared the figures, and wrote the original draft. ES supervised the study and reviewed and edited the manuscript. YY conceived and supervised the study, acquired funding, and reviewed and edited the manuscript. All authors read and approved the final manuscript.

---

**Supplementary Table S1.**
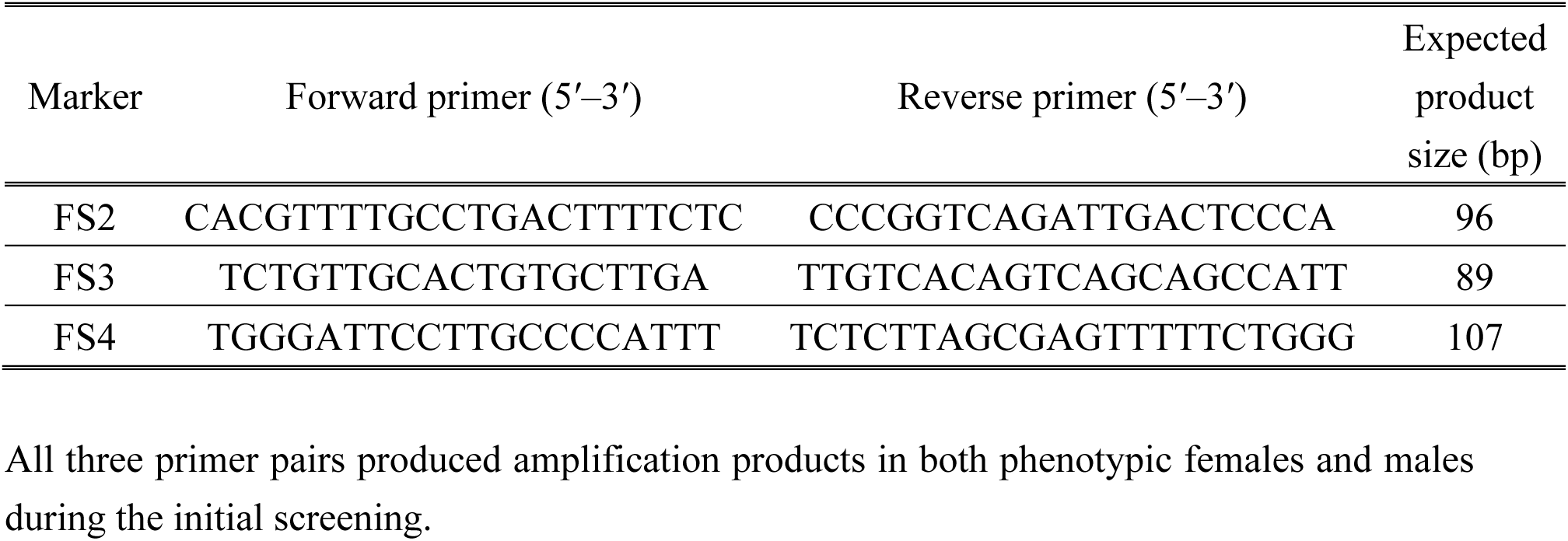
Primer sequences and PCR-screening results for three candidate female-associated markers that were excluded from further analysis.

**Supplementary Table S2.**
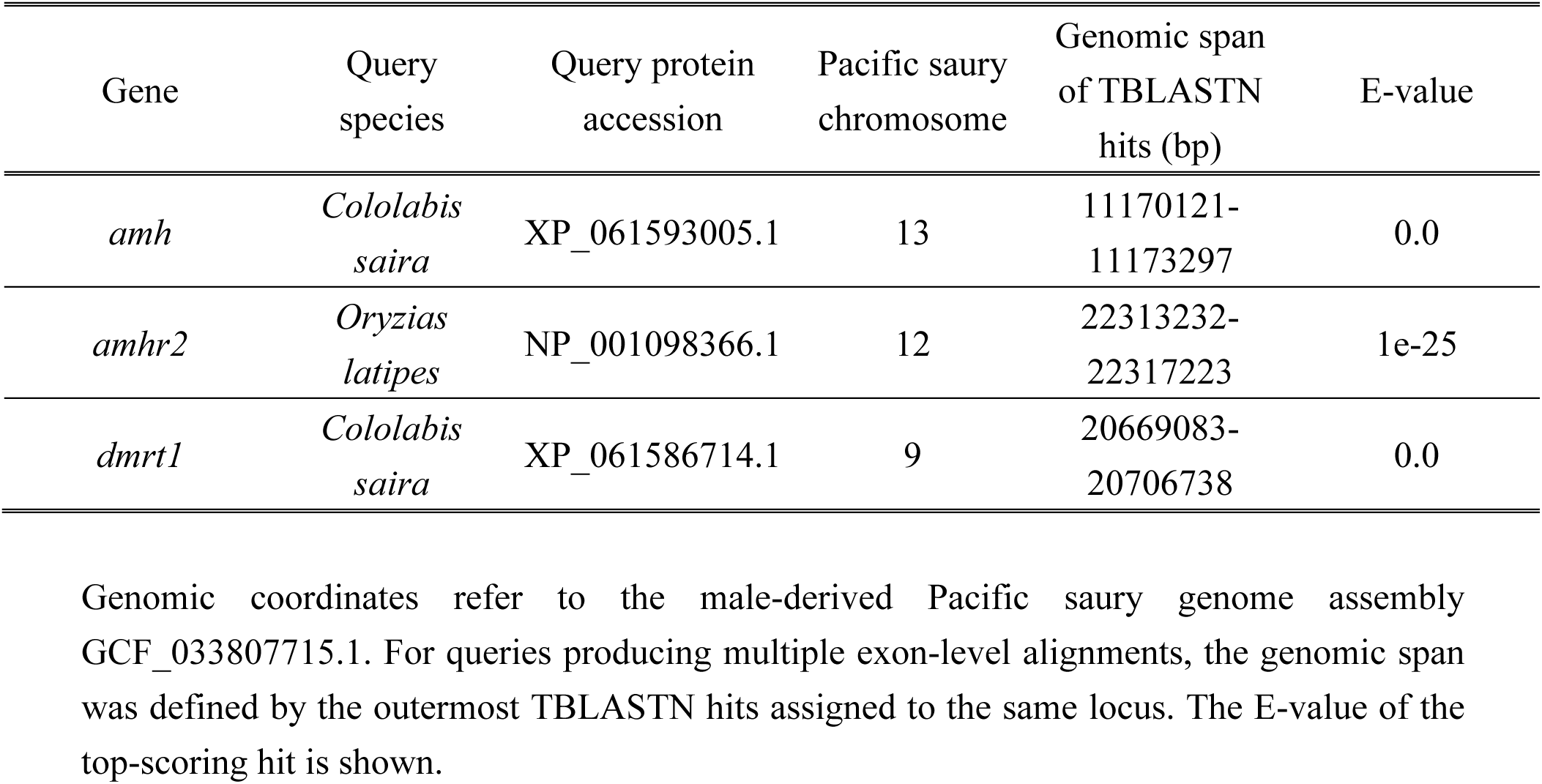
Putative genomic locations of sex-determination-related gene homologs in the Pacific saury genome identified by TBLASTN searches.

## Notes

### Competing Interest Statement

The authors have declared no competing interest.

